# *Drosophila* sperm carry a dietary sterol that increases offspring development

**DOI:** 10.64898/2026.06.02.729550

**Authors:** David Flores-Benitez, Anna Taubenberger, Michal Grzybek, Marko Brankatschk, Klaus Reinhardt

## Abstract

Diet affects male fitness directly and across generations. The current research focus is on dietary proteins and carbohydrates. Dietary lipids (DL), which are incorporated into cell membranes and affect signaling, were neglected or analyzed at pathological high-fat diet amounts. How fitness variation arises among males consuming healthy amounts of DL remains unknown, as do the fitness parameters being affected, including paternal effects. Contrasting mammal- vs. plant-derived DL, we measure mating rate, sperm count, sperm and seminal fluid quality, fertility and offspring growth in *Drosophila melanogaster*. We show that i) mammal-derived DL increased male fitness in isolation and competition, but without changing sperm numbers. ii) We discovered that lipid microcarriers in the seminal fluid fused with, and directly provided DL to sperm, changing sperm quality. iii) Dietary cholesterol, precursor of the growth hormone ecdysone, was incorporated into sperm, vectored to the zygote and improved offspring development and survival. This duality of *Drosophila* sperm function – fertilization and provision of growth hormone precursor – is an elegant, adaptive solution to prevent rival males from exploiting paternal provision. Direct offspring benefits vectored by sperm offer additional and alternative explanations for the evolution of sperm competition, sperm gigantism, and of anisogamy, the driver of sexual selection.

## Introduction

Diet crucially affects male reproduction, including all male (in)fertility parameters, including hormone levels, testicular traits, sperm counts, and semen quality^1,2^ (e.g., ^3–11^). Diet also affects the clinically and evolutionarily relevant health of the resulting offspring^12–16^, and therefore, have large transgenerational significance. Previous research in humans has focused on broad-spectrum diet differences (“Western” vs. “Mediterranean” diet) or used pathological sugar or dietary lipid (DL) quantities (e.g., high-fat diet)^12,13,17–19^. Previous research on animals concentrated on dietary proteins and carbohydrates^20–23^.

However, fitness variation remains largely unexplained among males consuming physiologically normal quantities of dietary lipids (DL) although the effects can be expected to be very large, and non-trivial, i.e. not merely governed by energy limitation^24^. DL can integrate into cellular membranes and cause substantial membrane remodeling^25^. Small alterations in the consumption of physiologically normal quantities of DL sources can strongly alter biophysical and functional properties of cells^26^, change physiological processes and organismal health and affect reproductive fitness^24,27^. For example, in humans dietary recommendations exist to improve sperm quality and numbers^28^ by keeping a specific diet, such as one rich in polyunsaturated fatty acids (PUFA), plant fats, or specific sterols^29,30^. In other species, similar observations are being made^24,31^.

Paternal diet effects on offspring are one form of paternal environmental effects^32^. Of these effects, two may be medically less relevant but have very large ecological and evolutionary implications^33,34^. First, the observation that non-genetic paternal provision can be ‘exploited’ by rival males suggests that, for example, a seminal fluid substance that enhances the growth of the offspring may be ‘exploited’ by a rival male fertilizing the eggs and so fathering the provisioned offspring. Second, paternal effects may act on females, which may then provide offspring. In this case, the paternal effect would, in fact, be the result of differential female allocation^32^. Therefore, “Identifying the proximate mechanism mediating paternal influence on offspring has become the ‘holy grail’ for paternal-effect studies“^32^.

Here we examine the three major pre- and post-mating male fitness components: sperm count, sperm quality, and zygote/offspring health focusing on DL. We aim to comprehensively test the idea that DL source affect fitness, to provide a mechanistic explanation for it and to examine how these DL may or may not be exploited by rival males, by partially excluding differential female allocation. We use a simple, general study system by feeding either animal- or plant-derived DL to *Drosophila melanogaster* males. The profiles of the two major lipid classes, sterols and fatty acids (FA) differ fundamentally between plant-and mammal-derived food^35,36^. The physiological mechanisms of lipid transport, storage and metabolism are functionally conserved across animals for both sterols and FA. For example, in humans and *Drosophila* the CD36 family regulates FA transport to the endoplasmic reticulum, or Niemann-Pick proteins and some ABC transporters^37,38^ regulate sterol absorption and transport. Sterols and FA produce the same biophysical membrane properties in humans as in *Drosophila* and affect male reproductive tissues^39^ although *Drosophila* relies on dietary sterols, while humans absorb dietary sterol species^40–42^. Both are auxotrophs for many PUFA species including omega-3 FA and get them from the diet to integrate them into their membranes. Below, we quantify DL effects on sperm numbers from the germ cell stage to storage in the female. We separate DL effects on sperm and seminal fluid, given that seminal fluid influences sperm movement, fertilizing capacity^43,44^, storage in the female^45^, and offspring health^5,46^ but that isolated effects of seminal DL are largely unknown for humans and *Drosophila*^43,44^. We reveal an unknown mechanism of DL delivery to sperm via seminal fluid. We quantify transgenerational DL effects and discover a mechanism how males provide direct benefits for their offspring (environmental paternal effect^32^). This benefit is directly tied to the paternal genome and so have low potential of being exploitable by rival males.

## Results

### Lipids of mammalian origin increase isolated male fertility

To quantify how DL impact male fertility, we used plant and mammalian lipidomes, which show little overlap in essential FA and sterol composition^35,36^. We fed males one of three isocaloric food sources: lipid-depleted food (LF), LF supplemented with either a lipid extract from either lard (Food*^Animal^*), or olive oil (Food*^Plant^*) (Table 1) (for the full treatment including sterol-only provision, see Supplementary material and supplementary fig. 1a). Food*^Animal^* and food*^Plant^* were not pathological (1% fat content), unlike high-fat diets (5-30% fat content) used in other studies^17^. Male flies that had DLs incorporated into cell membranes^47^ (Methods) were provided with ten virgin females (5-7 days old, reared on corn-based fly culture media) for 24h (supplementary fig. 1b). Males*^Plant^*and males*^Animal^* fertilized a similar number of females, and more than the males on lipid-free food (fig. 1a). However, males*^Animal^*produced more offspring than males^Plant^ (fig. 1a, supplementary fig. 1c-e). Counting the numbers of mature sperm stored in the seminal vesicles (SV) of males, we found low sperm numbers in SV of males^LF^, compared to DL-supplemented males, demonstrating that sperm production requires DL (fig. 1b). However, the number of non-differentiated cyst stem cells in the apical tip of the testis^48^ (fig. 1c) differed very little between males*^Animal^* and males*^Plant^* (fig. 1d) showing DL source did not influence stem cell proliferation rate. Our data also reject the possibilities that sperm of male*^Plant^*or males*^Animal^* (henceforth sperm*^Plant^* and sperm*^Animal^*) had higher success in being transferred to females (fig. 1e) or accepted or stored by them 24h after the start of mating (ASM) (fig. 1f). These data exclude sperm count as an explanation for variation in offspring number.

**Figure 1:**
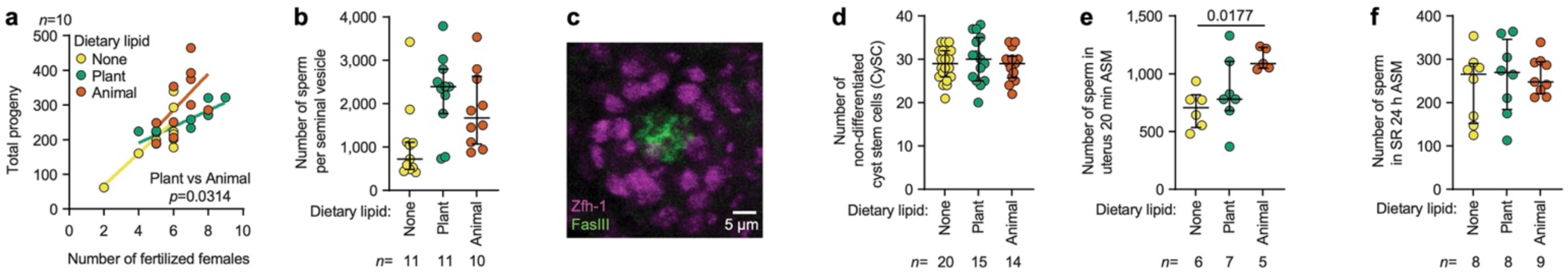
Males produce more offspring when kept on healthy amounts of animal than on plant lipids. Males were fed either lipid-free food (yellow, control) or lipid-free food supplemented with plant-derived (green) or animal-derived (brown) lipids. **a**, Males kept on animal lipids produced more offspring in total, and per female fertilized, than control males, and males kept on plant-derived lipid. **b**, The number of sperm produced and stored by males in the seminal vesicle was lower in males on the animal- than the plant-derived lipid diet. **c**,**d** Dietary lipids did not significantly change the number of non-differentiated cyst stem cells (Zfh-1 positive -magenta) located around the hub (FasIII positive -green) in the testis. **e**,**f**, Numbers of sperm counted in the uterus 20 min after the start of mating (ASM) (**e**) and the seminal receptacle (SR) 24 h ASM of individual females (**f**) did not differ between lipid treatments. n denotes the number of animals examined. The diet treatment also included a planned sterol-only supplementation and plant- and animal-derived lipids are not analyzed separately. For full statistical analysis see supplementary fig. 1.

**Table 1.**
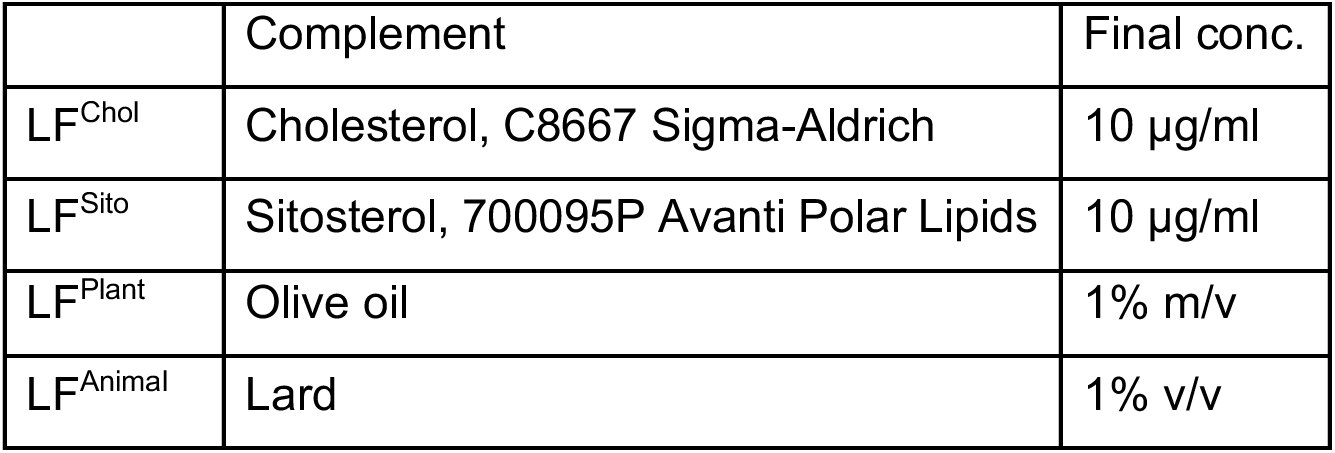
Composition and final concentrations of lipids in the food sources complementing lipid-free (LF) food used in this experiment.

|  | Complement | Final conc. |
| --- | --- | --- |
| LF<sup>Chol</sup> | Cholesterol, C8667 Sigma-Aldrich | 10 µg/ml |
| LF<sup>Sito</sup> | Sitosterol, 700095P Avanti Polar Lipids | 10 µg/ml |
| LF<sup>Plant</sup> | Olive oil | 1% m/v |
| LF<sup>Animal</sup> | Lard | 1% v/v |

Previous reports emphasized the role of sterols in sperm maturation and male fertility^39^. Isocaloric LF supplemented with only an animal-derived sterol, cholesterol (food*^Chol^*), or a plant-derived sterol, sitosterol (food*^Sito^*) rescued sperm numbers produced, transferred, and stored as much as food*^Animal^* and food*^Plant^* did (supplementary fig. 1h-k).

We next explored whether differences in sperm quality and function may explain DL-modulated fertility differences.

### Lipids of mammalian origin increase competitive male fertility

Multiple mating by females establishes the opportunity for postcopulatory sexual selection favoring males whose sperm is preferentially employed in fertilizations^49^. Thus, we focused on sperm competition offense and evaluated whether males under distinctive DL treatments produced ejaculates with different competing efficacy against sperm from a previous mating stored in the female. The paternity share of treatment males overall was highest in males^animal^ but showed few differences in paternity share (P2) with male DL treatment (fig. 2a). Males^Animal^ sired the highest absolute number of offspring under sperm competition (fig. 2b) but was not different from males^Plant^. Thus, the type of DL has a small influence on the ability of sperm to compete with sperm from other males.

**Figure 2:**
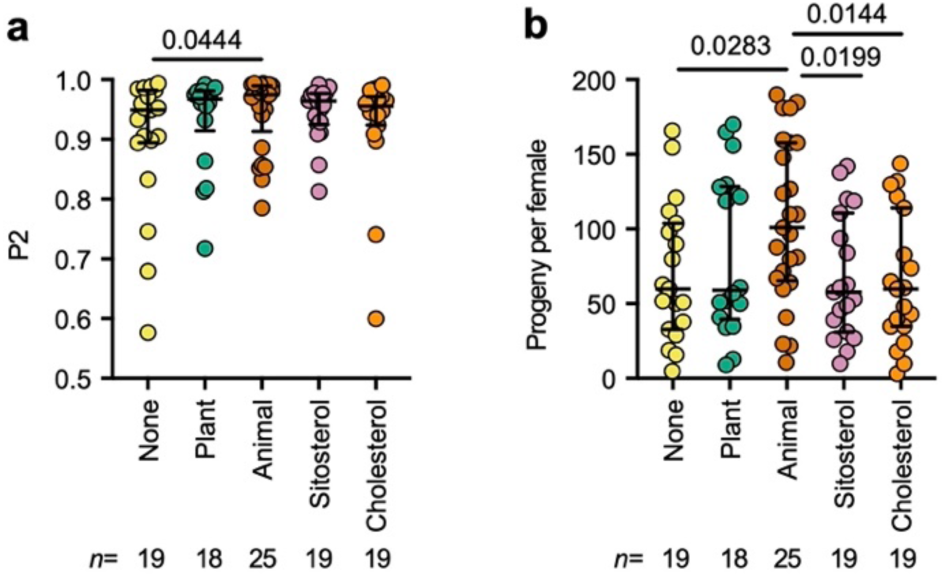
Males fed with animal-derived lipids have a small competitive fertility gain. Males fed a lipid-free food supplemented with different lipid sources were subjected to a sperm offense competition assay. **a**, Males fed animal-derived lipids sired a bigger P2 when compared to males fed a lipid-free medium. Kruskal-Wallis test, H(4)=4.618, p=0.3288. The p-value correspond to the FDR post-hoc test comparisons. **b**, Total number of adult offspring produced by females mated to males fed with the indicated DL types. Kruskal-Wallis test, H(4)=8.834, p=0.0654. The p-values correspond to the FDR post-hoc test comparisons between all groups. n denotes the number of individual matings examined.

### Sperm morphology and motility

Sperm morphological traits are key determinants of male fertility. Sperm*^Plant^* were ∼2% longer than sperm*^Animal^*(fig. 3a) but did not differ in tail diameter (fig. 3b,c). We also explored membrane tension by gauging the apparent Young’s modulus of sperm tails (fig. 3b,d). We found that *Drosophila* sperm are among the stiffest known cell types (values in the Megapascals range, fig. 3d), but found no DL effect on the stiffness of sperm tails (ANOVA p=0.4517, F(2,28)=0.8178).

**Figure 3:**
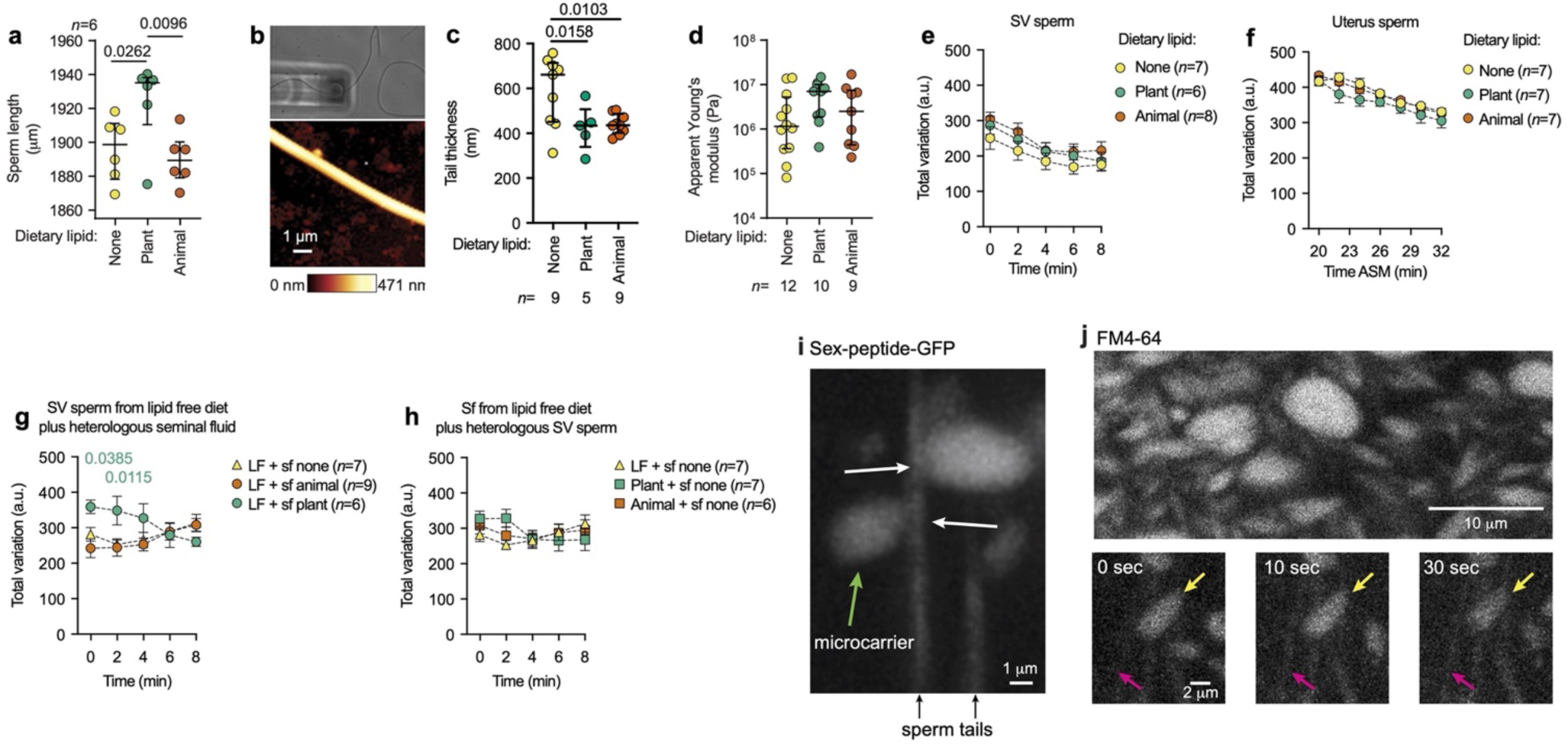
The sperm traits and motility change in the presence of dietary lipids. **a**, Sperm length of males kept on lipid-free food (yellow) and DL^Plant^ or DL^Animal^ (ANOVA p=0.0212, F(2,15)=5.041). Atomic force microscopy probing the sperm surface (**b**) showed thicker tails in control males than DL-treated males (ANOVA p=0.0145, F(2,20)=5.271) but no difference with the source of DLs provided (**c**). **d,** There was no difference in the apparent Young’s modulus (ANOVA p=0.4517, F(2,28)=0.8178). **e**, Seminal vesicle (SV) sperm were released in PBS, and immediately video recorded (see methods for details). Sperm motility was calculated as the total variation in pixel intensity over the 20 sec recordings at 50 frames per sec made every 2 min. **f**, Sperm from the uterus were released in PBS 20 min ASM, and immediately video recorded. **g**,**h**, Motility of sperm isolated from males kept on lipid-free food exposed to different Sf (**g**), and sperm isolated from males fed with either LF^Plant^ or LF^Animal^ and exposed to Sf from males kept on lipid-free food (**h**). Statistical significance in (**e**-**h**) was evaluated with a mixed-effects model followed by a FDR post-test. The colors for p-values correspond to the FDR post-hoc test comparisons between LF^Plant^ vs LF males, respectively. Only p-values < 0.05 are shown. Note that data in **g**,**h**, are separated for graphical clarity, but statistical analysis were performed using the whole data set together. Graphs show mean±SEM and the number of samples is indicated (n). **i**, Representative image showing fluorescent lipid microcarriers attached to a sperm tail; white arrows point to contact points (see supplementary movie 4). **j**, Top: representative image of the microcarriers loaded with FM4-64 inside the accessory glands. Bottom: frames from a time-lapse recording (supplementary movie 5) show a microcarrier attached to a sperm tail that becomes distinguishable over time as the microcarrier delivers its lipid content to the tail.

Sperm motility is a factor influencing male fertility^50^. We, therefore, next examined this important sperm quality parameter. Using DIC microscopy and high-speed time-lapse recordings (supplementary movie 1), we measured changes in pixel intensity caused by sperm tail beating and calculated the total variation of pixel intensity change over time (see Methods). When sperm were released from seminal vesicles (SV), all sperm types had similar motility (fig. 3e), suggesting that DL had no effect on sperm axoneme organization. Also, no motility differences between food treatments were observed in sperm extracted from the uterus 20 min after ASM (fig. 3f), yet uterus sperm had higher motility than SV sperm (fig. 3e vs 3f). We predicted that the interaction between seminal fluid (Sf) and sperm during mating would increase sperm motility and therefore tested whether Sf from males^Animal^ (Sf^Animal^) increases sperm motility more than Sf from males^Plant^ (Sf^Plant^). When we exposed sperm^LF^ to Sf^Plant^, sperm motility was initially higher than sperm^LF^ treated with Sf^LF^ or Sf^Animal^ (fig. 3g). However, after 6 min of incubation, the motility trends appear to invert, with sperm^LF^ treated with sf^LF^ or sf^Animal^ having higher motilities at the end of the recording than the beginning. Treatment of sperm^Plant^ or sperm^Animal^ with sf^LF^ showed no differences in motility response (fig. 3h). These data suggest that DLs determine the capacity of the seminal fluid to activate sperm motility.

Thus, the morphology and biophysical properties of sperm in the testes and the seminal vesicles were protected from the (small) compositional variation in DL type.

### Seminal fluid microcarriers deliver dietary lipids to sperm

Sf contains lipids organized in large containers called microcarriers^51^, among other components. The Sf lipid microcarriers disappeared rapidly (<90 s) in the presence of sperm but not in their absence (supplementary movie 2). We speculated that the microcarriers interact with sperm directly. We then used males whose Sf lipid microcarriers were labeled with the Sex peptide-GFP (SP-GFP)^52^ and followed the interaction of labeled microcarriers with non-fluorescent wild-type sperm (supplementary fig. 2a). We observed that Sf lipid microcarriers instantly associated with sperm cells (fig. 3i) and, once attached to the sperm tail, rapidly disappeared over time (supplementary movies 3 and 4). As predicted, sperm became visible only after fusion with microcarriers, when SP-GFP had associated to sperm tails. The cellular mechanisms of association and evacuation are unknown. However, we predicted that lipids in the microcarriers would be delivered to the sperm. We tested this by exposing unlabeled sperm to microcarriers labeled with the lipophilic probe FM4-64. Live-imaging showed that the FM4-64 probe appeared in the sperm after the binding of the microcarriers to the sperm tail (fig. 3j, supplementary movie 5), supporting the hypothesis that sperm receive lipids directly from the Sf microcarriers.

### Male dietary lipid type does not affect sperm metabolism

Sperm motility might be affected either via an altered energy metabolism or via cell membrane composition)^53^. To investigate any Sf-related alteration of sperm energy metabolism, we measured rates of oxidative phosphorylation (OXPHOS) relative to glycolysis. FLIM with time-correlated single photon count (TCSPC) allowed us to measure the autofluorescence of the metabolic cofactors nicotinamide adenine dinucleotide (phosphate) (NAD(P)H) and flavin adenine dinucleotide (FAD) in their free and protein-bound proportions. We calculated the FLIM-redox ratio of sperm (supplementary fig. 3e)^54,55^. Because OXPHOS consumes the NAD(P)H-bound fraction (NAD(P)H-a2% - i.e., that with a long lifetime, τ_2_=2.3-3.0 ns) and produces the FAD-bound fraction (FAD-a1% - i.e., that with a short lifetime, τ_1_=0.1-0.3 ns), the NAD(P)H-a2%:FAD-a1% ratio (FLIM-redox ratio) reflects the relative contribution of glycolysis and OXPHOS to the cell (sperm) redox state^54,55^. The metabolic profile of the stem cell compartment in the apical tip of the testis differed strongly from that of mature sperm stored in the seminal vesicles, and from sperm stored in the female seminal receptacle 40 min ASM (supplementary fig. 3a). Compared to the cells within the apical tip of the testis (which includes the germline stem cells), mature SV sperm stayed at a very low level of OXPHOS, and only glycolysis increased (supplementary fig. 3a). Sperm found in the female 40 min ASM (supplementary fig. 3a) had much higher OXPHOS levels than the mature SV sperm. These results show that within the female reproductive tract, sperm increased energy production via OXPHOS. However, the metabolic activity of sperm did not differ between sperm*^Plant^* and sperm*^Animal^* (supplementary fig. 3b-k) and continued to be similar 72 and 96h ASM in the female (supplementary fig. 3l-u). These results show that the DL source effects on male fertility and offspring success (fig. 1) do not act via DL effects on spermatogenesis and energy metabolism.

### Sperm cells carry dietary sterols that increase offspring developmental success

We found that after contact with Sf microcarriers, motility is affected by changes in plasma membrane organization, suggesting that DL-induced membrane properties caused motility differences. Yet, motility differences do not promote competitive interactions *per se* because food^Animal^ did not improve competitive over isolated fertility (fig. 2), suggesting that paternal DLs affect offspring number in other ways. Two candidates are the well-documented female-manipulating function of Sf that could lead to differential investment by females^56^ the other would be a direct transfer of Sf factors to the offspring. Such direct transfer would in our system only be possible via sperm. We first focused on the cholesterol, the main sterol of food^Animal^. We excluded female sources of cholesterol for offspring by feeding a cholesterol-free diet to females and mated them to either males*^Plant^* or males*^Animal^* (fig. 4a). We first confirmed previous data that the large *Drosophila melanogaster* sperm^57,58^, (0.1% of egg volume) enter the egg in totality^59,60^ (supplementary fig. 4g), we used mass spectrometry analysis of eggs fertilized ten days after mating. Zygotes from males*^Animal^* - but not males*^Plant^* - contained cholesterol (fig. 4b,c, supplementary fig. 4h-j). These data represent unequivocal evidence that sperm have a dual function in *Drosophila melanogaster* – fertilization and sterol delivery. Note that *Drosophila melanogaster* can feed on cholesterol in the wild^61^. Note also that sperm become degraded within three hours^62^ in the egg, in an endocytic pathway that recycles all membranes and tail components^63^, and remainders of which were detected in the midgut of the offspring^64^.

**Figure 4:**
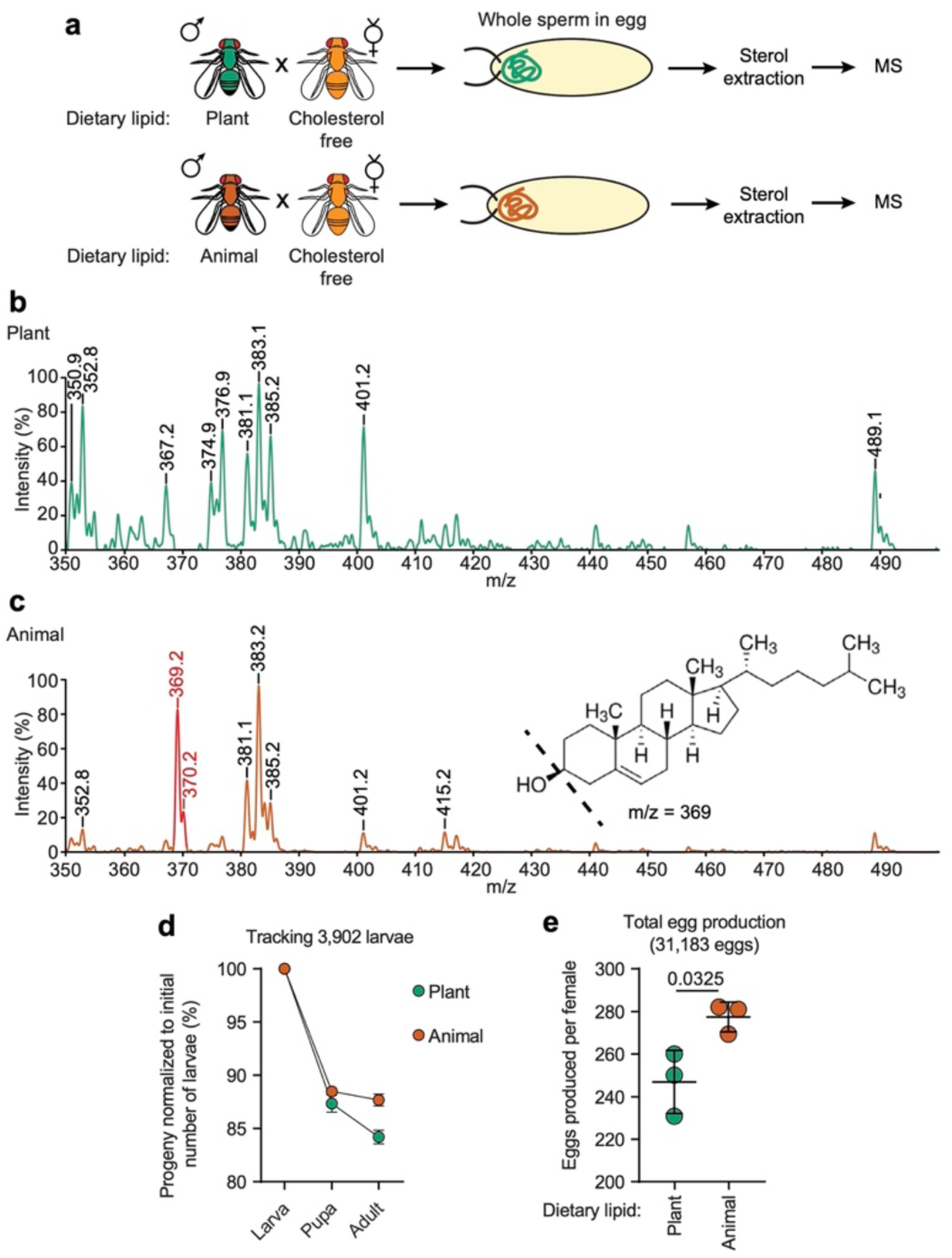
Sperm transport paternal sterols and modulate offspring developmental success. **a**, Males fed with DL^Plant^ or DL^Animal^ were mated to virgin females kept on cholesterol-free food. Eggs produced on day 10 ASM show specific mass spectrometry profiles (**b,c**) for cholesterol only in eggs fertilized by sperm^Animal^ (**c**) and not by sperm^Plant^ (**b**), - see peaks in red. Consult supplementary fig. 4h-j for the MS profiles of sterol standards. **d**, The development of the progeny from males kept on lipid-defined foods varied accordingly: more larvae pupate successfully and more adults eclosed from larvae that hatched from eggs fertilized by sperm^Animal^ than from those fertilized by sperm^Plant^. Graph show mean±SEM and the number of samples is 3 for each food. **e**, Females resulting from these treatments then differed in a further 10% in the number of eggs they laid (**e**, t=6.166, df=4), revealing a strong transgenerational signal. Graph show mean±SD.

Recent findings indicated that paternal diet^3,4^, especially high-fat lipid treatment^12,13,17–19^, can additionally act transgenerationally and affect offspring development and fertility. If these findings apply to the healthy amounts of DL used in our study and to sterol transmission, male treatment differences are predicted to result in hormone-based differences in offspring development. In *Drosophila melanogaster*, cholesterol is converted into 20-hydroxy-ecdysone (20HE), the essential hormone for larval growth and development into adults. Therefore, more precisely, the differences in offspring development should be seen in late development because early developmental failures may be caused by genomic damages of sperm. When examining the development of hatched larvae, we indeed found that offspring of males*^Animal^* had both higher pupation and higher adult eclosion rates than males*^Plant^* (fig. 4d, and supplementary fig. 4). These observations suggest that the sterol delivery to the offspring is adaptive in *Drosophila melanogaster*.

To assess the effect of sterols alone, we fed males with food supplemented with only mammalian-derived cholesterol or only the phytosterol sitosterol. Mostly, males*^Plant^* and males*^Animal^*males sired a similar number of offspring as males*^Sito^* and males*^Chol^*, respectively (supplementary fig. 1d), supporting the view the sperm-transmitted factor might be a sterol. Moreover, in *D. melanogaster*, the conversion of cholesterol into 20HE is known to be a thousand-fold more efficient than the conversion of phytosterols into Makisteron, the less potent 20HE equivalent^65,66^. This predicts, and our observation confirms this prediction, that males*^Sito^* produced fewer progeny than males*^Chol^* (fig. 1d,e). We conclude that *Drosophila melanogaster* males transfer dietary cholesterol to the offspring, thereby improving their development and survival.

We were not able to clearly decipher the role of differential female investment. We found that females produced about 10% more eggs after mating to males*^Animal^* than after mating to males*^Plant^*(fig. 4e), suggesting that the well-known fecundity-manipulating effect is related to DL source.

## Discussion

Our experiments using food supplemented with healthy amounts of dietary FA and sterols showed that sperm numbers, sperm quality and offspring success were all affected by the source of DL eaten by males. Our results indicate that such DL effects on offspring are based on different mechanisms than those recently reported for pathological high-fat diets ^17^. Our results not only concern healthy amounts of realistic food types, they also represent a novel way of direct paternal offspring provision^32^. The sterol provision shown here represents a bridging of the soma-germline division. Another important aspect of the paternal environment effect that we demonstrate is its tight coupling to the paternal genome. This tight coupling of the benefit to the offspring genome largely prevents an exploitation by other males.

Our evidence in the sterol auxotroph *Drosophila* that sterol transmission from father to offspring occurs via sperm has several strong and testable implications in a wide range of research disciplines. i) We predict that sperm-transmitted sterols may generally be an important resource for offspring. Whereas sterols are abundant in oocytes, they are esterified and less readily available than free sterols sexually transmitted by sperm. ii) We predict that sperm-transmitted male contributions may be phylogenetically widespread, because sperm tails enter the oocyte and are digested in other insects and *C. elegans* ^67^, in humans ^68^ and other mammals ^69–72^ and in fish ^73^. iii) At an ecological level, we expand the known DL flows through food webs ^24,27^, by two other routes that are as important for cell biologists as for ecologists: DL transmission from males to females, and across generations. iv) Sterol uptake adaptively differs with temperature in the wild ^47^, it is rapid, and may affect temperature-dependent responses (*10*). Male strategies to forage for fitness-increasing sterols can therefore be predicted to vary with temperature. Speculatively, environment-dependent female choice may evolve for those males that have loaded their sperm with the right sterol. Finally, vi) evolutionary models that rest on the assumption that males contribute little more than genetic material to the offspring may produce different outcomes than models taking into account direct male contributions. In addition to the above mentioned mate choice, direct male benefits that have low potential to be exploited by rival males may drive sperm gigantism and anisogamy. Direct male benefits may also affect sperm competition although in our model system the dietary alteration of sperm motility via seminal fluid did not. Direct and genome-coupled male offspring provision may produce similar effects as sperm epigenetics.

## Acknowledgments

We acknowledge the support of the Light Microscopy Facility of the Technology Platform of the Center for Molecular and Cellular Bioengineering of the TU Dresden. S. Broschk, C. Froschauer, A. Garside for food preparation, fly maintenance and excellent technical help, J. Pöschl for counting SV sperm, and N. Baboli for technical assistance in FLIM imaging. M. Garlovsky, R. Kraut, and S. Wigby for comments on the manuscript, M. Fedorova, Ü. Coskun and participants of three meetings to discuss aspects of our findings (Biology of Spermatozoa 2023, Nutritional Physiology section of International Congress of Entomology 2024, Insect Reproductive Molecules 2024). The fly stock expressing the Sp-GFP was a kind donation from the Clive Wilson lab. The study was funded by contributions of the DFG (RE 1666/9-1) to KR and FOR2682/TP1 to MB.

Contributions: experiments: DFB, AT, MB; mass spectrometry: MG; study design, conception and manuscript: DFB, KR and MB.

## Supplementary figures

**Supplementary fig. 1:**
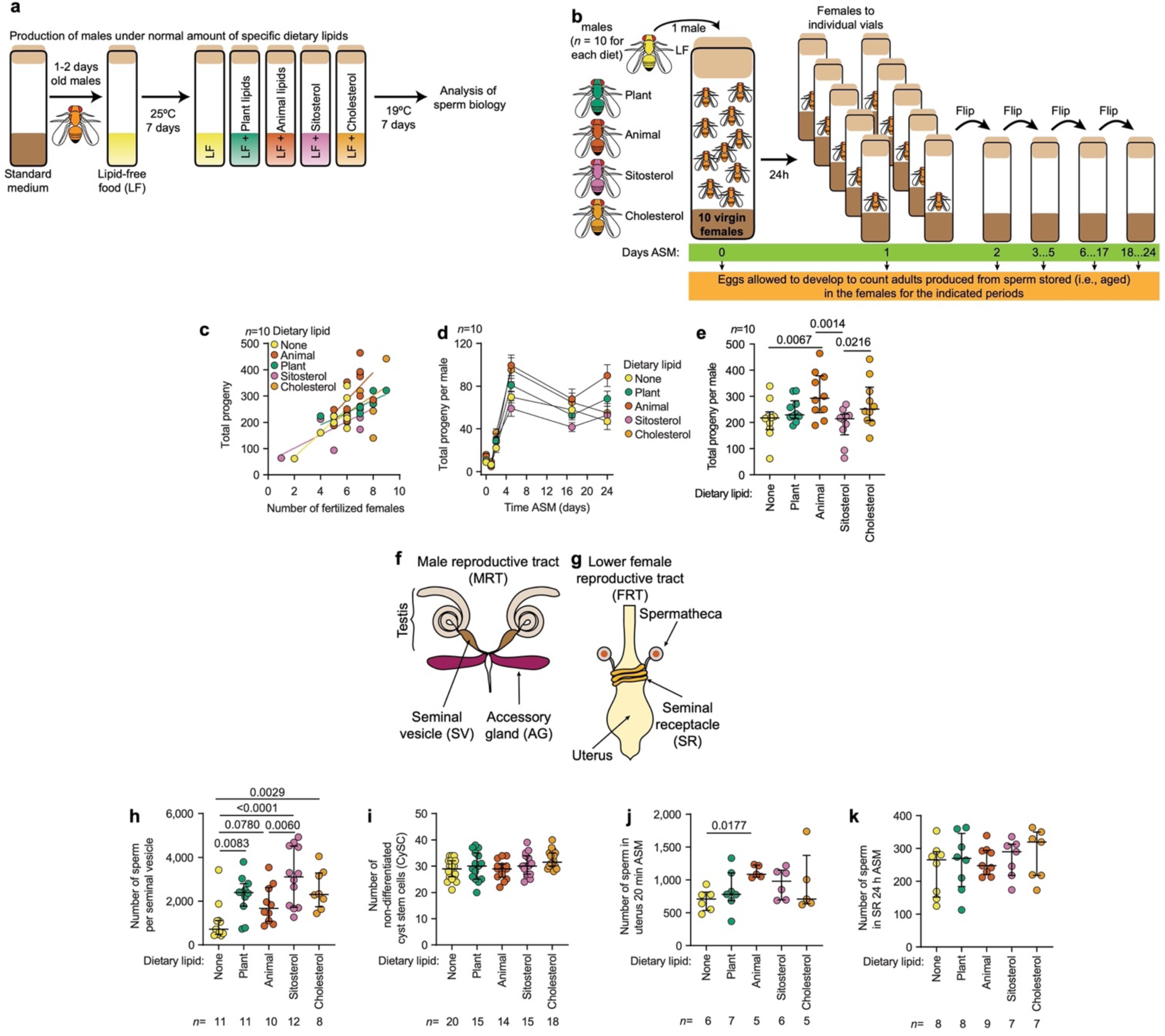
Males produce more offspring when kept on animal than on plant lipids. **a**, Males fed a lipid-free food supplemented with different lipid sources were subjected to fertility analysis. **b**, Experimental approach (see details in methods) to collect and count all progeny produced from single males mated to 10 females (n=10 males per food treatment). **c,d,e,** Total number of adult offspring produced by males in relation to the females fertilized (**c**) (Slope comparison (FDR-adjusted): None-Plant p=0.1509, None-Animal p=0.2757, None-Sitosterol p=0.2757, None-Cholesterol p=0.2757, Plant-Animal p=0.0314, Plant-Sitosterol p=0.5267, Plant-Cholesterol p=0.0314, Animal-Sitosterol p=0.0314, Animal-Cholesterol p=0.9420, Sitosterol-Cholesterol p=0.0314), and over time after the start of mating (ASM) (**d**)**. e**, Total progeny numbers produced by females mated to males fed with the indicated DL types (p=0.0330, one-way ANOVA, F(4, 45)=2.883). Reproductive tract of Drosophila melanogaster males (**f**) and females **g**, showing the location of sperm collection. **h,** Number of sperm males had stored in their sperm vesicle (premating) differed with diet (p=0.0003, F(4,47)=6.384). **i**, Dietary lipids did not significantly change the number of non-differentiated cyst stem cells (Zfh-1 positive -magenta) located around the hub (FasIII positive -green) in the testis (p=0.0517, one-way ANOVA, F(4,77)=2.468). **j**,**k**, No overall differences in numbers of sperm counted in the uterus 20 min after the start of mating (ASM) (**j**) (one-way ANOVA (p=0.8067, F(4, 34)=0.4007), and the seminal receptacle (SR) 24 h ASM of individual females (**k**) (FDR post-hoc tests following a one-way ANOVA (p=0.1647, F(4,24)=1.786)), n denotes the number of females. Plots show median, interquartile range (IQR), 75th percentile + 1.5 IQR, 25th percentile - 1.5 IQR.

**Supplementary fig. 2:**
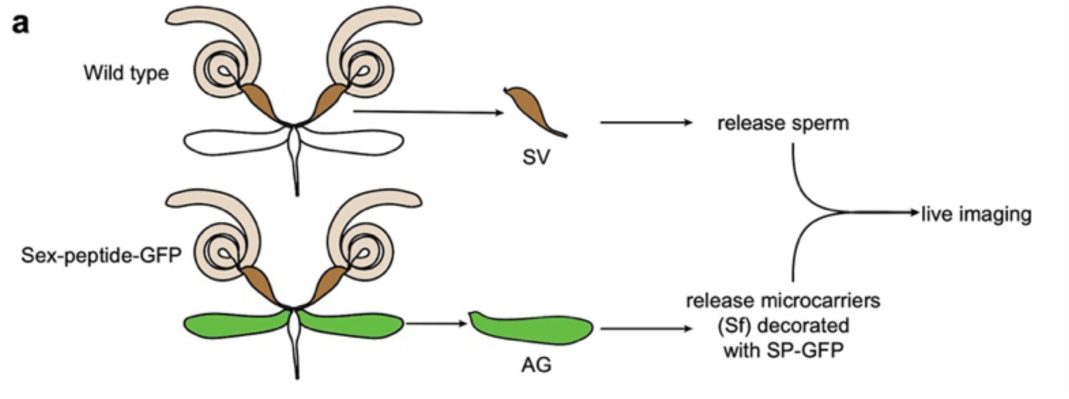
Strategy to combine sperm and seminal fluid from heterologous males. **a**, Experimental strategy to assess the interaction of Sf microcarriers with sperm in vitro

**Supplementary fig. 3:**
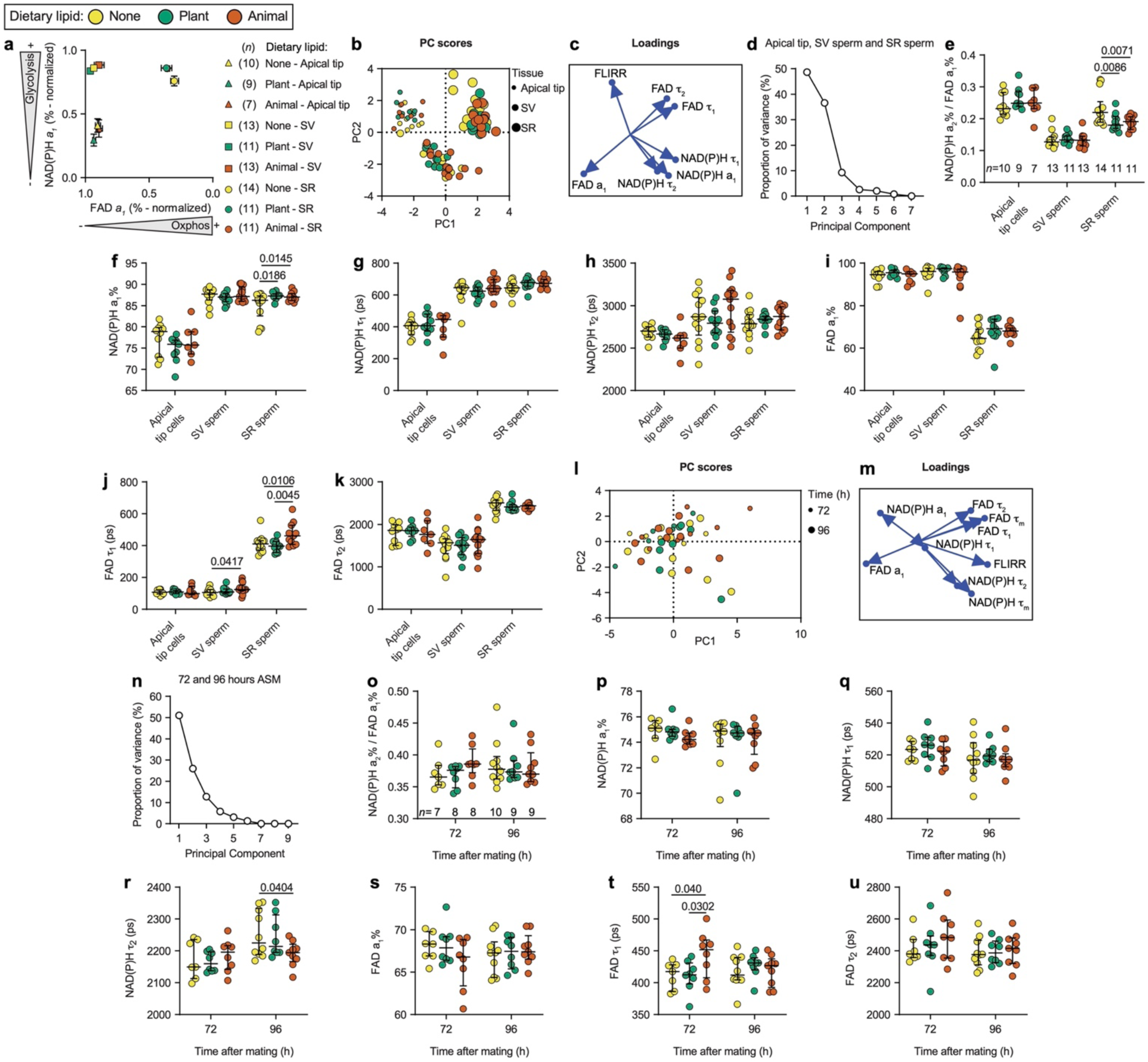
Metabolic activity of SV sperm and of sperm stored in females for 72h or 96h is not dependent on DLs consumed by males before mating. **a**, Sperm metabolic state evaluated by FLIM for normalized glycolysis (from NAD(P)H a1%) vs. normalized OXPHOS (from FAD a1%) shows strong separation between apical cells and SV sperm (sperm maturation), and between premating (SV) and postmating (SR sperm 40 min ASM) sperm. Data points are mean±SEM and (n) shows the number of replicates. No statistically significant difference was found between diet treatments within the different locations. **b-d**, PCA plot, PCA loadings and PCA proportions of variance from the metabolic state measured by FLIM of cells within the apical tip of the testis, SV sperm, and SR sperm 40 min ASM from males fed with LF, LF^Plant^, or LF^Animal^. **e**, Fluorescence-lifetime redox ratio (FLIRR) of cells within the testis apical tip, SV sperm, and SR sperm 40 min ASM from males fed with LF, LF^Plant^, or LF^Animal^. Plots show median, interquartile distance (IQR), 75th percentile + 1.5 IQR, 25th percentile - 1.5 IQR. An angular transformation was performed to percentage values before statistical analysis, and p-values were calculated using FDR post-hoc tests following a one-way ANOVA (SR sperm p=0.0083, F(2,33)=5.565). FLIM parameters were (**f**) NAD(P)H a1% (one-way ANOVA: SR sperm p=0.0195, F(2,33)=4.444), (**g**) NAD(P)H τ_1_, (**h**) NAD(P)H τ_2_, (**i**) FAD a1%, (**j**) FAD τ_1_ (one-way ANOVA: SV sperm p=0.1192, F(2,34)=2.266; SR sperm p=0.0088, F(2,33)=5.474), (**k**) FAD τ_2_. The number of samples (n) indicated in **e** applies also to **f-k**. **l-n**, PCA plot, PCA loadings and PCA proportions of variance from the metabolic state measured by FLIM of SR sperm 72 h and 96 h ASM from males fed with LF, LF^Plant^, or LF^Animal^. **o**, FLIRR of SR sperm 72 h and 96 h ASM from males fed with LF, LF^Plant^, or LF^Animal^. Calculated FLIM parameters are (**p**) NAD(P)H a1%, (**q**) NAD(P)H τ_1_, (**r**) NAD(P)H τ_2_ (one-way ANOVA: SR sperm 96 h ASM p=0.0898, F(2,24)=2.669), (**s**) FAD a1%, (**t**) FAD τ_1_ (one-way ANOVA: SR sperm 72 h ASM p=0.0510, F(2,20)=3.467), (**u**) FAD τ_2_. The number of samples (n) indicated in **o** applies also to **p-u**. Plots show median, interquartile distance (IQR), 75th percentile + 1.5 IQR, 25th percentile - 1.5 IQR. An angular transformation was performed to percentage values before statistical analysis.

**Supplementary fig. 4:**
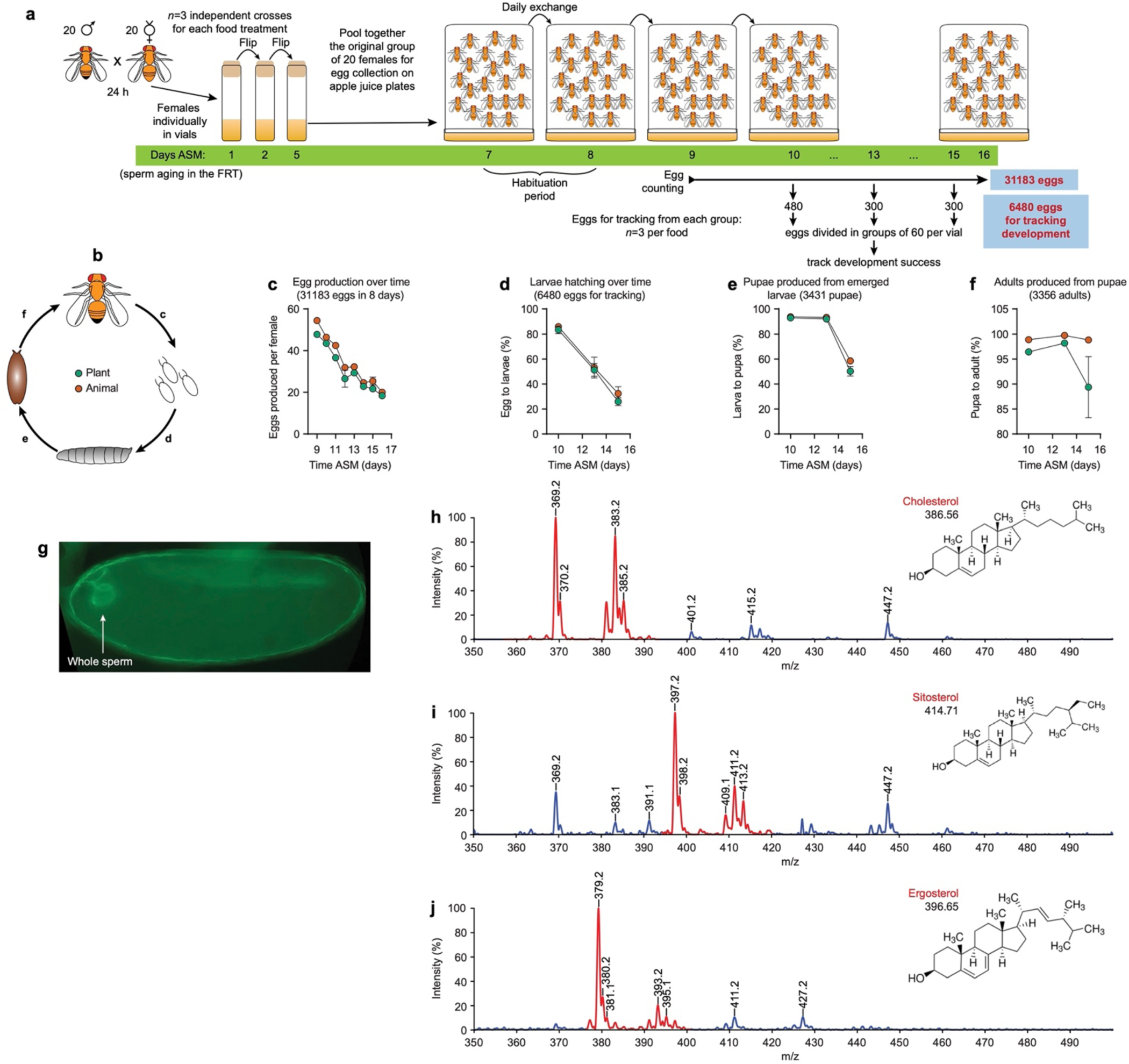
Developmental success of offspring sired. **a**, Experimental strategy to assess the developmental success of progeny from males kept on lipid-defined foods **b**, Drosophila melanogaster life cycle. **c**, Number of eggs produced by females. **d**, Percentage of larvae that developed successfully from eggs. **e**, Percentage of pupae from larvae hatched. **f**, Percentage of pupae able to eclose. Mean±SEM. n=3 independent crosses for each food. **g**, image of a recently (<1h) fertilized Drosophila egg with the sperm still visible, sperm is expressing Don Juan-GFP. Mass spectrometry profiles of (mammal-derived) cholesterol (**h**), (plant-derived) sitosterol (**i**), and (yeast-derived) ergosterol (**j**).

## Supplementary videos

**Movie S1**. High-speed time-lapse recordings of sperm released from the uterus 20 min ASM.

**Movie S2**. Seminal fluid (Sf) microcarriers in the presence of sperm (left) disappear in a few seconds as sperm motility increases. The microcarriers alone in PBS (right) remain visible during the same period of time.

**Movie S3**. Microcarrier (marked with Sex-peptide-GFP) attachment to sperm tails occurs promptly upon contact.

**Movie S4**. Microcarriers transfer their content to the sperm. Upon attachment, the GFP intensity on the microcarriers (blue and magenta dots) decreases rapidly while the GFP intensity remains relatively constant on the contiguous tail regions (yellow and orange dots).

**Movie S5**. Microcarriers transfer their lipid content to the sperm. Microcarriers were loaded with FM4-64 before adding them to unlabeled sperm. The microcarriers disappear rapidly while FM4-64 becomes detectable in the sperm tails.

## Materials and Methods

### *Drosophila* husbandry and dietary treatments

Canton S flies (Bloomington Stock Center 64349) were grown at 18-20°C on the standard food in our lab (1% malt extract, 8002-48-0 Carl Roth GmbH + Co. KG; 0.5% peptone ex soya, 91079-46-8 Carl Roth GmbH + Co. KG; 0.75% dry baking yeast Kaufland GmbH & Co. KG; 0.75% agar-agar, 9002-18-0 Carl Roth GmbH + Co. KG; 2% sucrose Netto ApS & Co. KG; 4% corn flour Kaufland GmbH & Co. KG; 0.3% nipagin, 99-76-3 Carl Roth GmbH & Co. KG). We produced males that integrate dietary lipids by maintaining 1-2 days old males on lipid-free food (LF: 10% D(+)-glucose, 50-99-7 Carl Roth GmbH + Co. KG; 10% yeast autolysate, 73145 Merk; 0.6 % agar-agar, 0.2% nipagin) at 25°C for 7 days; afterward, the males were transferred to LF complemented with specific lipids (Table 1) and kept at 19°C for 7 days. This protocol efficiently integrates dietary lipids in the *Drosophila* lipidome^47^. The corresponding lipid extracts from olive oil and lard were extracted after FOLCH^74^. Lipid fractions and sterols were added to warm LF food to evaporate the organic solvents

### Male fertility

For testing male fertility, we counted the number of females laying fertilized eggs and the total number of adult progeny sired by each male. For this, single males (14-15 days old) kept in the dietary treatment described above were placed individually with 10 virgin females (5-7 days old) and allowed to mate overnight at 19°C in vials with standard food. The females were separated into individual vials the following day and regularly checked for females laying unfertilized eggs. Females laying fertilized eggs were flipped 5 times to fresh food over 22 days. Vials were maintained at 20-22°C.

### Sperm competition assay

We collected virgin males and females from the *ebony* (*e*) line (Bloomington stock #497) and kept them in separate vials (20 individuals per vial) for 3-5 days. On day 1, couples of *e* females and males (1×1) were placed (without anesthetizing) in single vials with fresh standard media, and 24h later, the males were discarded. 40 individual crosses were set for each DL treatment to be tested. Unfertilized females identified by the lack of progeny after 48 h were discarded. Fertilized females were flipped to vials with fresh standard media on days 3, 5, and 7. On day 8, single CS males (14-15 days old) kept in the dietary treatment described above were placed individually with the fertilized *e* females, pairs were left to mate overnight, and males were discarded the next day. The *e* females were then flipped to vials with fresh standard media on days 9, 10, 11, 12, and 15. Vials were maintained at 20-22°C. To evaluate the proportion of progeny sired by each male, we score the number of adult *e* progeny (sired by the first male) and adult progeny with a wild-type appearance (sired by the second “lipid” male) emerging from each vial. The proportion of progeny sired by the second male (P2) was calculated using progeny numbers from vial 8 onward. We only scored vials that produced progeny from each mating, because that is where sperm mixing indeed occurred.

### Egg laying and development tracking

For testing success at each developmental stage, we used males (14-15 days old) in the dietary treatment described above. The males were put in groups of 20 together with 20 virgin females (5-7 days old) in standard food vials and allowed to mate overnight at 19°C. Three independent crosses were set for each food treatment. The females were separated into individual vials the following day, flipped to fresh-food vials on days 2 and 5 ASM, and kept at 24°C. All females were successfully fertilized, as observed by larvae present in all vials.

On day 7 ASM, the 20 females from each group were pooled together again in egg collection chambers (i.e., three independent cohorts per food treatment). Eggs were collected at 24°C on apple juice agar plates (2.6% M/V agar-agar Cat. 5210.3 ROTH, 25% V/V apple juice Fruchtstern Netto ApS & Co. KG, 2.5% M/V sucrose brand Südzucker AG Mannheim Germany, 1.5 mg/mL nipagin Cat. 3646.4 ROTH). The plates were exchanged daily. After a habituation period of 2 days, we counted laid eggs daily (a total of 31,183 eggs from day 9 to 16 ASM). All females stayed alive for the collection period, except one female that died on day 9 ASM in one replicate of the LF^Animal^. For this replicate, the number of eggs per female was calculated according to the remaining individuals for the corresponding days left.

Eggs collected on days 10, 13, and 15 ASM were used for tracking development. A total of 480 eggs (day 10 ASM) or 300 (days 13 and 15 ASM) from each cross (that is a total of 6480 eggs for all cohorts) were manually transferred into vials of fresh standard media as groups of 60 eggs per vial. All vials were kept at 24°C. To obtain the eclosion rate, we recorded the number of eggs that did not eclose the following day. At this point, it was also possible to count the numbers of eggs that were fertilized but did not eclose by the presence/absence of cuticles inside the eggs. Larvae were allowed to develop at 24°C, and we counted the numbers of pupae (a total of 3,431 pupae) and the numbers of imagoes emerging afterward (a total of 3,356 adults).

### Cell counting and sperm length

Non-differentiated cyst stem cells (CyCSs) were identified by antibodies against Zfh-1 and by their location next to hub cells identified by antibodies against Fasciclin-III (FasIII). Whole testes from males (14-15 days old) kept on the dietary treatments described above were dissected in ice-cold PBS and fixed at room temperature (R.T.) in 4% formaldehyde for 30 min, then washed 3 times with PBST (0.1 % Triton-X in PBS), permeabilized with 0.3% sodium deoxycholate (D6750, Sigma-Aldrich) in PBT for 30 min at R.T., and blocked overnight with 5% normal goat serum at 4°C. Incubation with primary antibodies was done in blocking solution overnight at 4°C: mouse anti-Fas-III (1:20, 7G10 Developmental Studies Hybridoma Bank) and rabbit anti-Zfh-1 (1:5000, kind gift from Ruth Lehmann)^75^. The samples were washed 3 times with PBST and then incubated with goat anti-mouse (1:500, Alexa Fluor 555, A32727 ThermoFisher) and goat anti-rabbit (1:500, Alexa FluorTM 488, A32731 ThermoFisher) in blocking solution at R.T. for 2 h. Samples were mounted in Vectashield (H-1000-10 Vector Laboratories), covered with a glass coverslip (1.5H, 0107052, Paul Marienfeld GmbH & Co. KG, Lauda-Koenigshofen, Germany), and visualized using a Zeiss LSM 780 AxioObserver single photon point scanning confocal system (Carl Zeiss Microscopy, Jena, Germany) with a Zeiss C-Apochromat 63x/1.20 W Korr M27 objective. For an experiment set (i.e., all samples from all conditions), all settings (laser power, detector sensitivity, pinhole, step size, and averaging) were kept constant. Images were analyzed using FIJI software. Brightness and contrast were changed equally for all channels and cohorts.

The number and length of sperm in the seminal vesicles (SV) were determined in males (14-15 days old) kept on the dietary treatments described above. One seminal vesicle per male was dissected in ice-cold PBS and transferred to 10 µl of PBS on a microscope slide. The SV was carefully opened with sharp tweezers, and sperm were allowed to spread. The excess PBS was carefully removed without drying the sample, 5 µL of mounting solution was added (50% glycerol, 5 µg/ml DAPI (D1306 Invitrogen) and 2% formaldehyde (Sigma F8775) in PBS) and covered with a cover slip (#1.5H). The samples were visualized using a Zeiss Axio Scan.Z1 slide scanner with a Zeiss Plan-Apochromat 20x/0.8 M27 objective. 16-bit images were captured with a pixel size of 0.325 µm x 0.325 µm. Image processing was performed using ZEN 3.0 (blue edition, Carl Zeiss Microscopy GmbH 2019), and the number and length of sperm (head + tail) were scored manually using FIJI software^76^.

To evaluate the number of sperm transferred after mating, males (14-15 days old) kept on the different dietary treatments were individually mated to virgin females 5-6 days old. Matings were timed, and males were discarded after mating was completed. For counting sperm in the uterus, females were frozen 20 min after the start of mating (ASM). For counting sperm in the seminal receptacle (SR) at 24h, 7 days, and 12 days ASM, the males were discarded after mating finished, and the females were kept in groups of 10 animals/vial in standard food at 20°C and then frozen at the corresponding times. For dissections, the females were thawed in ice-cold PBS, the lower female reproductive tract (FRT) was dissected in ice-cold PBS, transferred to 5 µl of PBS on a microscope slide, and then covered with 10 µL of mounting solution (50% glycerol, DAPI 5 µg/ml and formaldehyde 2% in PBS). Samples were visualized using a Zeiss LSM 780 AxioObserver single photon point scanning confocal system (Carl Zeiss Microscopy, Jena, Germany) with a Zeiss Plan-Apochromat 20x/0.8 M27 objective. The number of sperm was scored manually using FIJI.

### Fluorescence lifetime imaging microscopy (FLIM)

Metabolic FLIM for measuring the lifetimes of nicotinamide adenine dinucleotide (phosphate) (NAD(P)H) and flavin adenine dinucleotide (FAD) was performed as previously reported, and all sample collections and recordings were randomized^55^. In detail, the male reproductive tract (excluding the ejaculatory bulb) of flies (14-15 days old) kept on the different dietary treatments were dissected in ice-cold PBS, transferred to 10 µl of PBS on a microscope slide prepared with a spacer (double-sided tape (05338 tesa), covered with a coverslip (#1.5H), and imaged immediately. Different sets of males were employed for FLIM measurements of cells within the apical tip of the testis (comprising the germline stem cells, spermatocytes, and supporting cells) and for the sperm within the SV. For the FLIM measurements of sperm stored in SR 40 min ASM, males (14-15 days old) kept under the specific dietary treatments were individually mated to virgin females 5-6 days old. Single matings were observed and timed. The male was discarded 40 min ASM, and the female reproductive tract was dissected in ice-cold PBS, transferred to 10 µl of PBS on a microscope slide prepared with a spacer (double-sided tape) covered with a coverslip (#1.5H), and imaged immediately. All FLIM data (imaging) was acquired at 20-22°C, and the internal-reference-function (IRF) was measured for each imaging day using a slide with urea crystals. Fluorescence lifetime data was collected using time-correlated single photon counting (TCPSC) FLIM with an AxioExaminer.Z1 (Carl Zeiss Microscopy, Jena, Germany) with a Zeiss LD C-Apochromat 40x/1.1 W Korr M27 objective, a Chameleon Ultra II two-photon titanium:sapphire laser (tunable range 690–1080 nm, 80 MHz repetition rate, 140 fs pulse, Coherent, USA) and two hybrid GaAsP photon detectors (HPM-100-40, Becker & Hickl GmbH, Berlin, Germany). NAD(P)H and FAD fluorescence was two-photon excited with 740 nm and 900 nm, respectively. Emitted light was split with a 505 nm beam splitter. A bandpass filter of 450/30 nm and 525/39 nm was used for NAD(P)H and FAD fluorescence recording, respectively. Images (340 frames, 256×256 pixel format, zoom 3, 147.3 s scanning time) were acquired with SPCM software version 9.80 (Becker&Hickl GmbH, Berlin, Germany). Fluorescence decays were corrected with the corresponding IRF before fitting. To extract the values for *1*_1_, *1*_2_, a1, and a2, NAD(P)H and FAD fluorescence decays were fitted to a two-component exponential model with a fixed scatter of 0 using SPCImage software 8.0 (Becker&Hickl GmbH, Berlin, Germany). For all samples, regions of interest (ROIs) were manually selected in images of NAD(P)H mean lifetimes *1*_m_ and then applied to the corresponding FAD images.

For the metabolic FLIM analysis of sperm in the SR at 72h and 96h ASM, males (14-15 days old) kept on specific dietary regimes were mated to virgin females (3-4 days old). Matings were observed, and males were discarded after the mating was finished. Females were maintained at 20°C in standard food until the time of dissection. The lower FRT was dissected in ice-cold PBS, transferred to 10 µl of PBS on a microscope slide prepared with a spacer, covered with a coverslip (#1.5), and imaged immediately using time-correlated single photon counting (TCPSC) FLIM with a Leica SP8 FALCON (Leica Microsystems, Wetzlar, Germany) with a Fluotar VISIR 25x/0.95 Water objective, a Tunable Ultrafast Laser (tunable range 680-1300 nm, 80 MHz repetition rate, InSight X3, Spectra-Physics, MKS Instruments Inc., USA). NAD(P)H and FAD fluorescence were two-photon excited with 740 nm and 900 nm, respectively, and detected using HyD detectors in the non-descanned (NDD) config.uration (NDD1 range 435-470 nm for NAD(P)H; NDD2 range 500-550 nm for FAD). Images (20 frame repetitions, 512×512 pixel format, zoom 3) were acquired with LAS X software 3.5.7.23225 (Leica Microsystems, Wetzlar, Germany). To extract the values for *1*_1_, *1*_2_, a1, and a2, NAD(P)H and FAD fluorescence decays were fitted to a bi-exponential reconvolution model using LAS X FLIM/FCS software version 3.5.6 (Leica Microsystems, Wetzlar, Germany). For all samples, the same regions of interest (ROIs) for NAD(P)H and FAD channels were manually selected in images of “Fast FLIM” mean lifetimes *1*_m_.

### Sterol mass spectrometry from thin layer chromatography plate

Canton S males prepared as described above were allowed to mate for 4h to 3-5 days old virgin Canton S females (10 males x 10 females in vials with standard media, 3 independent crosses per food treatment). Males were discarded the next day, and the females were kept in vials with standard media at 24°C and transferred to fresh vials every other day. On day 7 ASM, the females were placed in egg collection chambers (i.e., three independent cohorts per food treatment). Eggs were collected at 24°C on apple juice agar plates (2.6% M/V agar-agar Cat. 5210.3 ROTH, 25% V/V apple juice Fruchtstern Netto ApS & Co. KG, 2.5% M/V sucrose brand Südzucker AG Mannheim Germany, 1.5 mg/mL nipagin Cat. 3646.4 ROTH). The plates were exchanged daily. On day 10 ASM, eggs were collected for 24 h. The following morning, 20 eggs per plate were transferred to an Eppendorf tube and put at −80°C till analysis. For the analysis, the samples were applied to Silica 60 HPTLC plate (Merck) and separated in a Heptane-Dimethyl ether-Acetic (70:30:1) acid running solvent. The plate was stained with primuline solution (0.005%) and visualized with a 380nm LED lamp. Areas corresponding to cholesterol were marked with pencil. TLC/MS analyses were performed using the Plate Express extraction device coupled to the expression CMS-L single quadrupole MS (Advion Scientific Interchim). This device incorporates an extraction head that forms a leak-tight seal on the surface of the TLC plate, which allows an extraction solvent to be delivered onto the TLC spot followed by a direct coupling to the inlet of the compact API mass spectrometer. The TLC plate extraction solvent/spray solvent was methanol for the elution of the analytes from the TLC plate. Positive APCI MS analyses with a m/z scan range 350-500 was recorded.

### Atomic force microscopy (AFM)

To measure individual sperm tails, SV from males (14-15 days old) kept in the dietary regimes described above were dissected in ice-cold PBS, transferred to a glass-bottom petri dish (FluoroDish FD35-100, World Precision Instruments) coated the day before with 50 μL poly-L-lysine (P4707, SIGMA) by overnight incubation at 37°C. Sperm were released by carefully opening the SV with sharp tweezers and allowed to attach to the PLL. Then, the dish was carefully filled with PBS and mounted onto the sample stage of a Nanowizard 4 (JPK Instruments/Bruker) setup combined with a Zeiss inverted light microscope (AxioObserverZ1). Qpbio cantilevers (Nanoworld, Switzerland) were calibrated before AFM quantitative imaging and force mapping using the thermal noise calibration procedure implemented in the SPM software (version 6.1.159). Using phase contrast microscopy, an area containing an immobilized and well-accessible sperm tail was selected, and the tip was lowered onto the surface. Then, a preselected rectangular region (typically 3µm x 5µm) was scanned using the quantitative imaging mode (resolution appr. 16nm per pixel, tip speed 50µm/sec, relative setpoint 0.5nN, z length 1µm). After finishing the scan, a smaller region was selected in the obtained image using the SPM software, and an array of force-distance curves was recorded on a rectangular grid (e.g., 2µm x 3µm, 50nm distance between points, relative setpoint 2.5 nN, z length 2µm, speed 5µm/sec). Force distance curves recorded on top of the sperm tails were analyzed using the JPK/Bruker data processing software (version 6.1.X), choosing the Hertz/Sneddon model for a conical indenter (half-angle 22.5°, assuming a Poisson ratio of 0.5). For analysis of tail dimensions, quantitative images were exported as grayscale 8-bit tiff images to FIJI, and tail heights and width were measured using grey scale (z) and x-y scales in multiple profiles (plot profile function) perpendicular to the tail axis (8 profiles per image). All measurements were conducted at room temperature (18-20°C).

### Sperm motility

We analyzed sperm motility in three different conditions from males (14-15 days old) kept on specific dietary treatments. (1) Mature sperm released from SV, (2) sperm released from uterus 20 min ASM, and (3) mature sperm released from SV but incubated with homo- or heterologous seminal fluid. All dissections were done in ice-cold PBS, avoiding squeezing the SV or uterus to avoid mechanical disturbance of the sperm by the tweezers. For (1), the SV was transferred to 5 µl of PBS on a microscope slide with a spacer. The sperm was released by tearing open the SV. For (2), males were individually mated to virgin females 5-6 days old. The male was discarded 20 min ASM, then the female reproductive tract was dissected in ice-cold PBS, transferred to 5 µl of PBS on a microscope slide with a spacer, and the sperm was released by carefully opening the uterus. For (3), the AGs and SV from the same male (homologous seminal fluid treatment) or from two males on different dietary regimes (heterologous seminal fluid treatment) were isolated and placed on a slide with 5 µl of PBS. Then, the AGs were open to release the Sf, and the sperm was released from the SV afterward.

Data collection was randomized, and all recordings started within 20 seconds after the sperm release. Otherwise, the sample was discarded. Recordings were done in phase contrast microscopy using an inverted Leica DM IRB with an N PLAN 10x/0,22 (Nr. 506018) objective and a Phantom Miro M310 high-speed camera (Ametek Vision Research). For each sample (i.e., each male), at least five recordings of 20 s spaced by 2 min each were acquired with the Phantom Camera Control software (Version 2.8.761.0, Vision Research Inc.) at a resolution of 384×288 pixels at 50 pps. Uncompressed .avi files were transformed into .tiff image sequences (of 1000 frames each) using FIJI. To assess sperm motility, we analyzed changes in pixel intensities resulting from the beating of sperm tails. We used a FIJI macro to randomly generate 30 ROIs of 2×2 pixels in the regions of the recording where sperm tails beating was visible. We then extracted, for each frame and ROI, the standard deviation (StdDev) using the measurement tools in FIJI. The StdDev values were further processed using the R package version 4.5.1; the change in pixel intensity was used as a proxy to estimate the intensity of tail beating, which we consider to be sperm motility. First, we calculated the difference in StdDev values between frames for each ROI in each recording. Then we calculated the sum of the absolute differences for each ROI in each recording. The median value of the 30 ROIs from one recording was then used as the “Total variation” value for that sample in that time point. The exact regions for a particular sample were analyzed in each of the recordings. All time points were considered and analyzed using a Mixed-effects model, followed by a multiple comparison test using the False Discovery Rate method of Benjamini and Hochberg run in GraphPad Prism 9.5.1 to evaluate statistical differences between the dietary treatments. Results are reported in fig. 3e-h as mean±SEM.

### Live imaging and FM4-64

To visualize the interaction of the seminal fluid microcarriers with sperm, we used males expressing the Sex-peptide tagged with GFP (w*; Sp-GFP, kindly donated by the Clive Wilson lab) and CS males 10 days old kept on standard medium. SVs were dissected as described in the sperm motility section above. The SVs were transferred to slides coated the day before with 50 µL poly-L-lysine (PLL, P4707, SIGMA) by overnight incubation at 37°C. The accessory glands (AGs) from SP-GFP males was dissected on ice-cod PBS and a single gland was transferred to the slide with the SV from CS males. Sperm were released by carefully opening the SV with sharp tweezers and allowed to attach to the PLL. Then, the AG was rupture to release the seminal fluid in a way that the seminal fluid (microcarriers) could bathe the sperm. The mix was cover with a coverslip (#1.5) and visualized immediately using a Zeiss LSM 780 AxioObserver single photon point scanning confocal system (Carl Zeiss Microscopy, Jena, Germany) with a Zeiss LSM 780 AxioObserver single photon point scanning confocal system (Carl Zeiss Microscopy, Jena, Germany), with a 488 nm excitation laser, and with a Zeiss C-Apochromat 40x/1.20 W Korr FCS M27 objective.

To visualize the interaction of sperm with the seminal fluid microcarriers loaded with FM4-64, we used CS males 10 days old kept on standard medium. The SV were dissected and placed on slides with PLL as described in the previous paragraph. The AGs were dissected on iced-cold PBS and then transferred to a 10 µL drop of FM4-64 10 µM in PBS and incubated for 60 sec at R.T. The AGs were then washed three times by serial transferring the glands into drops of 150 µL fresh PBS. The AGs were then transferred to the slide with the SV, the sperm was released and allowed to attach the PLL, and then the seminal fluid was released by opening the AGs. The mix was cover with a coverslip (#1.5) and visualized immediately using a Zeiss LSM 880 AxioObserver single photon point scanning confocal system (Carl Zeiss Microscopy, Jena, Germany) with a Zeiss LSM 880 AxioObserver single photon point scanning confocal system (Carl Zeiss Microscopy, Jena, Germany), with a 488 nm excitation laser, and with a Zeiss C-Apochromat 40x/1.20 W Korr FCS M27 objective.

### Figures preparation

All figures were made using Adobe Illustrator version 28.5. Microscopy images were processed with FIJI software. Lifetime images were generated using LAS X FLIM/FCS software version 3.5.6 (Leica Microsystems, Wetzlar, Germany).

### Statistics

All data (and the data for fig. 1A and supplementary fig. 1c) was analyzed using GraphPad Prism 9.5.1 (713). An angular transformation was applied to percentages before statistical evaluation. Gaussian distribution was evaluated using in-built normality tests in GraphPad Prism. Data fitting a normal distribution was analyzed by an ordinary one-way ANOVA. When data failed to pass the normality tests, statistical differences were evaluated by the Kruskal-Wallis test. A *p*-value of less than 0.05 was considered statistically significant. A multiple comparison test using the False Discovery Rate (FDR) by Benjamini and Hochberg was used to evaluate statistical differences between dietary treatment cohorts. The error bars indicate the 75^th^ percentile + 1.5 IQR (unless this value is greater than the largest value, in which case the whisker stops at the largest) and the 25^th^ percentile – 1.5 IQR (unless this value is smaller than the lowest value, in which case the whisker stops at the lowest value). The regression analysis of the data in fig. 1a and supplementary fig. 1c was done with R using the emmeans package (Lenth R (2025). emmeans: Estimated Marginal Means, aka Least-Squares Means. R package version 1.11.2-00002, https://rvlenth.github.io/emmeans/).

## Notes

### Competing Interest Statement

The authors have declared no competing interest.

## References

1. Leisegang, K., and Dutta, S. (2021). Do lifestyle practices impede male fertility? Andrologia 53. 10.1111/and.13595.

2. Crean, A.J., Afrin, S., Niranjan, H., Pulpitel, T.J., Ahmad, G., Senior, A.M., Freire, T., Mackay, F., Nobrega, M.A., Barrès, R., et al. (2023). Male reproductive traits are differentially affected by dietary macronutrient balance but unrelated to adiposity. Nat Commun 14, 2566. 10.1038/s41467-023-38314-x.

3. Zeender, V., Pfammatter, S., Roschitzki, B., Dorus, S., and Lüpold, S. (2023). Genotype-by-environment interactions influence the composition of the *Drosophila* seminal proteome. Proc. R. Soc. B. 290, 20231313. 10.1098/rspb.2023.1313.

4. Solon-Biet, S.M., Walters, K.A., Simanainen, U.K., McMahon, A.C., Ruohonen, K., Ballard, J.W.O., Raubenheimer, D., Handelsman, D.J., Le Couteur, D.G., and Simpson, S.J. (2015). Macronutrient balance, reproductive function, and lifespan in aging mice. Proc. Natl. Acad. Sci. U.S.A. 112, 3481–3486. 10.1073/pnas.1422041112.

5. Reinhardt, K., Dobler, R., and Abbott, J. (2015). An Ecology of Sperm: Sperm Diversification by Natural Selection. Annu. Rev. Ecol. Evol. Syst. 46, 435–459. 10.1146/annurev-ecolsys-120213-091611.

6. Morimoto, J., and Wigby, S. (2016). Differential effects of male nutrient balance on pre- and post-copulatory traits, and consequences for female reproduction in Drosophila melanogaster. Sci Rep 6, 27673. 10.1038/srep27673.

7. Salas-Huetos, A., Bulló, M., and Salas-Salvadó, J. (2017). Dietary patterns, foods and nutrients in male fertility parameters and fecundability: a systematic review of observational studies. Hum Reprod Update 23, 371–389. 10.1093/humupd/dmx006.

8. Macartney, E.L., Crean, A.J., Nakagawa, S., and Bonduriansky, R. (2019). Effects of nutrient limitation on sperm and seminal fluid: a systematic review and meta-analysis. Biological Reviews 94, 1722–1739. 10.1111/brv.12524.

9. Guo, R., and Reinhardt, K. (2020). Dietary polyunsaturated fatty acids affect volume and metabolism of *Drosophila melanogaster* sperm. J of Evolutionary Biology 33, 544–550. 10.1111/jeb.13591.

10. Schjenken, J.E., Moldenhauer, L.M., Sharkey, D.J., Chan, H.Y., Chin, P.Y., Fullston, T., McPherson, N.O., and Robertson, S.A. (2021). High-fat Diet Alters Male Seminal Plasma Composition to Impair Female Immune Adaptation for Pregnancy in Mice. Endocrinology 162, bqab123. 10.1210/endocr/bqab123.

11. Rodak, K., and Kratz, E.M. (2023). PUFAs and Their Derivatives as Emerging Players in Diagnostics and Treatment of Male Fertility Disorders. Pharmaceuticals 16, 723. 10.3390/ph16050723.

12. Nätt, D., Kugelberg, U., Casas, E., Nedstrand, E., Zalavary, S., Henriksson, P., Nijm, C., Jäderquist, J., Sandborg, J., Flinke, E., et al. (2019). Human sperm displays rapid responses to diet. PLoS Biol 17, e3000559. 10.1371/journal.pbio.3000559.

13. Öst, A., Lempradl, A., Casas, E., Weigert, M., Tiko, T., Deniz, M., Pantano, L., Boenisch, U., Itskov, P.M., Stoeckius, M., et al. (2014). Paternal Diet Defines Offspring Chromatin State and Intergenerational Obesity. Cell 159, 1352–1364. 10.1016/j.cell.2014.11.005.

14. Rando, O.J. (2012). Daddy Issues: Paternal Effects on Phenotype. Cell 151, 702–708. 10.1016/j.cell.2012.10.020.

15. Watkins, A.J., Dias, I., Tsuro, H., Allen, D., Emes, R.D., Moreton, J., Wilson, R., Ingram, R.J.M., and Sinclair, K.D. (2018). Paternal diet programs offspring health through sperm- and seminal plasma-specific pathways in mice. Proc. Natl. Acad. Sci. U.S.A. 115, 10064–10069. 10.1073/pnas.1806333115.

16. Morgan, H.L., Paganopoulou, P., Akhtar, S., Urquhart, N., Philomin, R., Dickinson, Y., and Watkins, A.J. (2020). Paternal diet impairs F1 and F2 offspring vascular function through sperm and seminal plasma specific mechanisms in mice. The Journal of Physiology 598, 699–715. 10.1113/JP278270.

17. Tomar, A., Gomez-Velazquez, M., Gerlini, R., Comas-Armangué, G., Makharadze, L., Kolbe, T., Boersma, A., Dahlhoff, M., Burgstaller, J.P., Lassi, M., et al. (2024). Epigenetic inheritance of diet-induced and sperm-borne mitochondrial RNAs. Nature 630, 720–727. 10.1038/s41586-024-07472-3.

18. Crisóstomo, L., Jarak, I., Rato, L.P., Raposo, J.F., Batterham, R.L., Oliveira, P.F., and Alves, M.G. (2021). Inheritable testicular metabolic memory of high-fat diet causes transgenerational sperm defects in mice. Sci Rep 11, 9444. 10.1038/s41598-021-88981-3.

19. Gómez-Elías, M.D., Rainero Cáceres, T.S., Giaccagli, M.M., Guazzone, V.A., Dalton, G.N., De Siervi, A., Cuasnicú, P.S., Cohen, D.J., and Da Ros, V.G. (2019). Association between high-fat diet feeding and male fertility in high reproductive performance mice. Sci Rep 9, 18546. 10.1038/s41598-019-54799-3.

20. Simpson, S.J., and Raubenheimer, D. (2012). The Nature of Nutrition: A Unifying Framework from Animal Adaptation to Human Obesity (Princeton University Press) 10.23943/princeton/9780691145655.001.0001.

21. Raubenheimer, D., and Simpson, S.J. (1997). Integrative models of nutrient balancing: application to insects and vertebrates. Nutr. Res. Rev. 10, 151–179. 10.1079/NRR19970009.

22. Lee, K.P., Simpson, S.J., Clissold, F.J., Brooks, R., Ballard, J.W.O., Taylor, P.W., Soran, N., and Raubenheimer, D. (2008). Lifespan and reproduction in *Drosophila*: New insights from nutritional geometry. Proc. Natl. Acad. Sci. U.S.A. 105, 2498–2503. 10.1073/pnas.0710787105.

23. Shingleton, A.W., Masandika, J.R., Thorsen, L.S., Zhu, Y., and Mirth, C.K. (2017). The sex-specific effects of diet quality versus quantity on morphology in *Drosophila melanogaster*. R. Soc. open sci. 4, 170375. 10.1098/rsos.170375.

24. Trautenberg, L.C., Brankatschk, M., Shevchenko, A., Wigby, S., and Reinhardt, K. (2022). Ecological lipidology. eLife 11, e79288. 10.7554/eLife.79288.

25. Levental, K.R., Malmberg, E., Symons, J.L., Fan, Y.-Y., Chapkin, R.S., Ernst, R., and Levental, I. (2020). Lipidomic and biophysical homeostasis of mammalian membranes counteracts dietary lipid perturbations to maintain cellular fitness. Nat Commun 11, 1339. 10.1038/s41467-020-15203-1.

26. Brankatschk, M., Gutmann, T., Knittelfelder, O., Palladini, A., Prince, E., Grzybek, M., Brankatschk, B., Shevchenko, A., Coskun, Ü., and Eaton, S. (2018). A Temperature-Dependent Switch in Feeding Preference Improves Drosophila Development and Survival in the Cold. Developmental Cell 46, 781–793.e4. 10.1016/j.devcel.2018.05.028.

27. Twining, C.W., Bernhardt, J.R., Derry, A.M., Hudson, C.M., Ishikawa, A., Kabeya, N., Kainz, M.J., Kitano, J., Kowarik, C., Ladd, S.N., et al. (2021). The evolutionary ecology of fatty-acid variation: Implications for consumer adaptation and diversification. Ecology Letters 24, 1709–1731. 10.1111/ele.13771.

28. Jensen, T.K., Heitmann, B.L., Jensen, M.B., Halldorsson, T.I., Andersson, A.-M., Skakkebæk, N.E., Joensen, U.N., Lauritsen, M.P., Christiansen, P., Dalgård, C., et al. (2013). High dietary intake of saturated fat is associated with reduced semen quality among 701 young Danish men from the general population. The American Journal of Clinical Nutrition 97, 411–418. 10.3945/ajcn.112.042432.

29. Esmaeili, V., Shahverdi, A.H., Moghadasian, M.H., and Alizadeh, A.R. (2015). Dietary fatty acids affect semen quality: a review. Andrology 3, 450–461. 10.1111/andr.12024.

30. Nassan, F.L., Chavarro, J.E., and Tanrikut, C. (2018). Diet and men’s fertility: does diet affect sperm quality? Fertility and Sterility 110, 570–577. 10.1016/j.fertnstert.2018.05.025.

31. Ugine, T.A., Krasnoff, S.B., Grebenok, R.J., Behmer, S.T., and Losey, J.E. (2019). Prey nutrient content creates omnivores out of predators. Ecology Letters 22, 275–283. 10.1111/ele.13186.

32. Crean, A.J., and Bonduriansky, R. (2014). What is a paternal effect? Trends in Ecology & Evolution 29, 554–559. 10.1016/j.tree.2014.07.009.

33. Bonduriansky, R., and Day, T. (2009). Nongenetic Inheritance and Its Evolutionary Implications. Annu. Rev. Ecol. Evol. Syst. 40, 103–125. 10.1146/annurev.ecolsys.39.110707.173441.

34. Bonduriansky, R., Mallet, M.A., Arbuthnott, D., Pawlowsky-Glahn, V., Egozcue, J.J., and Rundle, H.D. (2015). Differential effects of genetic vs. environmental quality in *Drosophila melanogaster* suggest multiple forms of condition dependence. Ecology Letters 18, 317–326. 10.1111/ele.12412.

35. Ahmad Nizar, N.N., Nazrim Marikkar, J.M., and Hashim, D.M. (2013). Differentiation of Lard, Chicken Fat, Beef Fat and Mutton Fat by GCMS and EA-IRMS Techniques. J. Oleo Sci. 62, 459–464. 10.5650/jos.62.459.

36. Cavanna, D., Hurkova, K., Džuman, Z., Serani, A., Serani, M., Dall’Asta, C., Tomaniova, M., Hajslova, J., and Suman, M. (2020). A Non-Targeted High-Resolution Mass Spectrometry Study for Extra Virgin Olive Oil Adulteration with Soft Refined Oils: Preliminary Findings from Two Different Laboratories. ACS Omega 5, 24169–24178. 10.1021/acsomega.0c00346.

37. Baldán, Á., Bojanic, D.D., and Edwards, P.A. (2009). The ABCs of sterol transport. Journal of Lipid Research 50, S80–S85. 10.1194/jlr.R800044-JLR200.

38. Plummer, A.M., Culbertson, A.T., and Liao, M. (2021). The ABCs of Sterol Transport. Annu. Rev. Physiol. 83, 153–181. 10.1146/annurev-physiol-031620-094944.

39. Keber, R., Rozman, D., and Horvat, S. (2013). Sterols in spermatogenesis and sperm maturation. Journal of Lipid Research 54, 20–33. 10.1194/jlr.R032326.

40. Ostlund, R.E., McGill, J.B., Zeng, C.-M., Covey, D.F., Stearns, J., Stenson, W.F., and Spilburg, C.A. (2002). Gastrointestinal absorption and plasma kinetics of soy Δ^5^-phytosterols and phytostanols in humans. American Journal of Physiology-Endocrinology and Metabolism 282, E911–E916. 10.1152/ajpendo.00328.2001.

41. Lafi, A.Sh.A., Tzar, M.N., Santhanam, J., and Huyop, F. (2024). Comparing ergosterol identification by HPLC with fungal serology in human sera. Heliyon 10, e38377. 10.1016/j.heliyon.2024.e38377.

42. Simonen, P., Öörni, K., Sinisalo, J., Strandberg, T.E., Wester, I., and Gylling, H. (2023). High cholesterol absorption: A risk factor of atherosclerotic cardiovascular diseases? Atherosclerosis 376, 53–62. 10.1016/j.atherosclerosis.2023.06.003.

43. Pitnick, S., Wolfner, M.F., and Dorus, S. (2020). Post-ejaculatory modifications to sperm (PEMS). Biological Reviews 95, 365–392. 10.1111/brv.12569.

44. McCullough, E.L., Whittington, E., Singh, A., Pitnick, S., Wolfner, M.F., and Dorus, S. (2022). The life history of *Drosophila* sperm involves molecular continuity between male and female reproductive tracts. Proc. Natl. Acad. Sci. U.S.A. 119, e2119899119. 10.1073/pnas.2119899119.

45. Avila, F.W., Sirot, L.K., LaFlamme, B.A., Rubinstein, C.D., and Wolfner, M.F. (2011). Insect Seminal Fluid Proteins: Identification and Function. Annu. Rev. Entomol. 56, 21–40. 10.1146/annurev-ento-120709-144823.

46. Bromfield, J.J., Schjenken, J.E., Chin, P.Y., Care, A.S., Jasper, M.J., and Robertson, S.A. (2014). Maternal tract factors contribute to paternal seminal fluid impact on metabolic phenotype in offspring. Proc. Natl. Acad. Sci. U.S.A. 111, 2200–2205. 10.1073/pnas.1305609111.

47. Knittelfelder, O., Prince, E., Sales, S., Fritzsche, E., Wöhner, T., Brankatschk, M., and Shevchenko, A. (2020). Sterols as dietary markers for Drosophila melanogaster. Biochimica et Biophysica Acta (BBA) - Molecular and Cell Biology of Lipids 1865, 158683. 10.1016/j.bbalip.2020.158683.

48. Zoller, R., and Schulz, C. (2012). The Drosophila cyst stem cell lineage: Partners behind the scenes? Spermatogenesis 2, 145–157. 10.4161/spmg.21380.

49. Fiumera, A.C., Dumont, B.L., and Clark, A.G. (2005). Sperm Competitive Ability in Drosophila melanogaster Associated With Variation in Male Reproductive Proteins. Genetics 169, 243–257. 10.1534/genetics.104.032870.

50. Yang, Y., and Lu, X. (2011). Drosophila Sperm Motility in the Reproductive Tract1. Biology of Reproduction 84, 1005–1015. 10.1095/biolreprod.110.088773.

51. Wainwright, S.M., Hopkins, B.R., Mendes, C.C., Sekar, A., Kroeger, B., Hellberg, J.E.E.U., Fan, S.-J., Pavey, A., Marie, P.P., Leiblich, A., et al. (2021). *Drosophila* Sex Peptide controls the assembly of lipid microcarriers in seminal fluid. Proc. Natl. Acad. Sci. U.S.A. 118, e2019622118. 10.1073/pnas.2019622118.

52. Villella, A., Peyre, J.-B., Aigaki, T., and Hall, J.C. (2006). Defective transfer of seminal-fluid materials during matings of semi-fertile fruitless mutants in Drosophila. J Comp Physiol A 192, 1253–1269. 10.1007/s00359-006-0154-1.

53. Serafini, S., and O’Flaherty, C. (2025). Novel insights into the lipid signalling in human spermatozoa. Human Reproduction 40, 1440–1451. 10.1093/humrep/deaf085.

54. Wallrabe, H., Svindrych, Z., Alam, S.R., Siller, K.H., Wang, T., Kashatus, D., Hu, S., and Periasamy, A. (2018). Segmented cell analyses to measure redox states of autofluorescent NAD(P)H, FAD & Trp in cancer cells by FLIM. Sci Rep 8, 79. 10.1038/s41598-017-18634-x.

55. Wetzker, C., and Reinhardt, K. (2019). Distinct metabolic profiles in Drosophila sperm and somatic tissues revealed by two-photon NAD(P)H and FAD autofluorescence lifetime imaging. Sci Rep 9, 19534. 10.1038/s41598-019-56067-w.

56. Avila, F.W., Sirot, L.K., LaFlamme, B.A., Rubinstein, C.D., and Wolfner, M.F. (2011). Insect Seminal Fluid Proteins: Identification and Function. Annual Review of Entomology 56, 21–40. 10.1146/annurev-ento-120709-144823.

57. Joly, D., Korol, A., and Nevo, E. (2004). Sperm Size Evolution in Drosophila: Inter- and Intraspecific Analysis. Genetica 120, 233–244. 10.1023/B:GENE.0000017644.63389.57.

58. Hill, A., and Eisenberg, D. (2016). The long and the short of it: new insights on sperm length help demystify the complexities of sexual selection. Asian J Androl 18, 902. 10.4103/1008-682X.185850.

59. Karr, T.L. (1991). Intracellular sperm/egg interactions in Drosophila: A three-dimensional structural analysis of a paternal product in the developing egg. Mechanisms of Development 34, 101–111. 10.1016/0925-4773(91)90047-A.

60. Loppin, B., and Karr, T.L. (2005). Molecular Genetics of Insect Fertilization. In Comprehensive Molecular Insect Science (Elsevier), pp. 213–236. 10.1016/B0-44-451924-6/00001-6.

61. Knittelfelder, O., Prince, E., Sales, S., Fritzsche, E., Wöhner, T., Brankatschk, M., and Shevchenko, A. (2020). Sterols as dietary markers for Drosophila melanogaster. Biochimica et Biophysica Acta (BBA) - Molecular and Cell Biology of Lipids 1865, 158683. 10.1016/j.bbalip.2020.158683.

62. Ben-Hur, S., Sernik, S., Afar, S., Kolpakova, A., Politi, Y., Gal, L., Florentin, A., Golani, O., Sivan, E., Dezorella, N., et al. (2024). Egg multivesicular bodies elicit an LC3-associated phagocytosis-like pathway to degrade paternal mitochondria after fertilization. Nat Commun 15, 5715. 10.1038/s41467-024-50041-5.

63. Politi, Y., Gal, L., Kalifa, Y., Ravid, L., Elazar, Z., and Arama, E. (2014). Paternal Mitochondrial Destruction after Fertilization Is Mediated by a Common Endocytic and Autophagic Pathway in Drosophila. Developmental Cell 29, 305–320. 10.1016/j.devcel.2014.04.005.

64. Pitnick, S., and Karr, T.L. (1998). Paternal products and by–products in Drosophila development. Proc. R. Soc. Lond. B 265, 821–826. 10.1098/rspb.1998.0366.

65. Lavrynenko, O., Rodenfels, J., Carvalho, M., Dye, N.A., Lafont, R., Eaton, S., and Shevchenko, A. (2015). The Ecdysteroidome of *Drosophila*: influence of diet and development. Development, dev.124982. 10.1242/dev.124982.

66. Kannangara, J.R., Mirth, C.K., and Warr, C.G. (2021). Regulation of ecdysone production in *Drosophila* by neuropeptides and peptide hormones. Open Biol. 11, 200373. 10.1098/rsob.200373.

67. Zhou, Q., Li, H., and Xue, D. (2011). Elimination of paternal mitochondria through the lysosomal degradation pathway in C. elegans. Cell Res 21, 1662–1669. 10.1038/cr.2011.182.

68. Wallace, D.C. (2007). Why Do We Still Have a Maternally Inherited Mitochondrial DNA? Insights from Evolutionary Medicine. Annu. Rev. Biochem. 76, 781–821. 10.1146/annurev.biochem.76.081205.150955.

69. Sutovsky, P., Moreno, R.D., Ramalho-Santos, J., Dominko, T., Simerly, C., and Schatten, G. (2000). Ubiquitinated Sperm Mitochondria, Selective Proteolysis, and the Regulation of Mitochondrial Inheritance in Mammalian Embryos1. Biology of Reproduction 63, 582–590. 10.1095/biolreprod63.2.582.

70. Sutovsky, P., McCauley, T.C., Sutovsky, M., and Day, B.N. (2003). Early Degradation of Paternal Mitochondria in Domestic Pig (Sus scrofa) Is Prevented by Selective Proteasomal Inhibitors Lactacystin and MG1321. Biology of Reproduction 68, 1793–1800. 10.1095/biolreprod.102.012799.

71. Sutovsky, P., Navara, C.S., and Schatten, G. (1996). Fate of the Sperm Mitochondria, and the Incorporation, Conversion, and Disassembly of the Sperm Tail Structures during Bovine Fertilization1. Biology of Reproduction 55, 1195–1205. 10.1095/biolreprod55.6.1195.

72. Song, W.-H., Yi, Y.-J., Sutovsky, M., Meyers, S., and Sutovsky, P. (2016). Autophagy and ubiquitin–proteasome system contribute to sperm mitophagy after mammalian fertilization. Proc. Natl. Acad. Sci. U.S.A. 113. 10.1073/pnas.1605844113.

73. Nishimura, Y., Yoshinari, T., Naruse, K., Yamada, T., Sumi, K., Mitani, H., Higashiyama, T., and Kuroiwa, T. (2006). Active digestion of sperm mitochondrial DNA in single living sperm revealed by optical tweezers. Proc. Natl. Acad. Sci. U.S.A. 103, 1382–1387. 10.1073/pnas.0506911103.

74. Eggers, L.F., and Schwudke, D. (2016). Liquid Extraction: Folch. In Encyclopedia of Lipidomics, M. R. Wenk, ed. (Springer Netherlands), pp. 1–6. 10.1007/978-94-007-7864-1_89-1.

75. Leatherman, J.L., and DiNardo, S. (2008). Zfh-1 Controls Somatic Stem Cell Self-Renewal in the Drosophila Testis and Nonautonomously Influences Germline Stem Cell Self-Renewal. Cell Stem Cell 3, 44–54. 10.1016/j.stem.2008.05.001.

76. Schindelin, J., Arganda-Carreras, I., Frise, E., Kaynig, V., Longair, M., Pietzsch, T., Preibisch, S., Rueden, C., Saalfeld, S., Schmid, B., et al. (2012). Fiji: an open-source platform for biological-image analysis. Nat Methods 9, 676–682. 10.1038/nmeth.2019.

